# High-throughput tuning of ovarian cancer spheroids for on-chip invasion assays

**DOI:** 10.1101/2021.11.24.469887

**Authors:** Changchong Chen, Yong He, Elliot Lopez, Franck Carreiras, Ayako Yamada, Marie-Claire Schanne-Klein, Ambroise Lambert, Yong Chen, Carole Aimé

## Abstract

We developed an invasion assay by using microfabricated culture devices. First, ovarian tumor spheroids were generated with a culture patch device consisting of an agarose membrane formed with a honeycomb microframe – the *patch* – and gelatin nanofiber backbone. By changing the dimensions of the honeycomb compartments we were able to control the number of cells and size of the spheroids. When the spheroids were placed on a patch coated with a thin membrane of fibrillary type I collagen, spheroid disruption was observed due to substrate induced cell migration. This process is straightforward and should be applicable to other cancer types, as well as assays under microfluidic conditions, thereby holding the potential for use in tumor modeling and anti-cancer drug development.

## 1. Introduction

Progress in cancer research relies partially on our ability to provide relevant *in vitro* models, which recapitulate important features of tumor growth, metastasis, and angiogenesis [1]. *In vitro* generation of tumor spheroids, for example, is a simple yet reliable approach for predicting tumor growth dynamics and microenvironmental dependencies [2,3]. Spheroids are formed by self-assembly of cells through different types of intercellular interactions, including cadherin- and integrin-mediated interactions, as well as involving the vitronectin/αv integrin adhesive system [4]. Ovarian cancer (OC) cells are likely to form spheroids, which seem to be the preferential mode of survival of floating OC cells, as well as the starting point for metastatic activity. The formation of such aggregates may contribute to protect them from the circulating microenvironment. Indeed, cells within spheroids have been shown to resist anoikis [5] and to be more prone to resist chemotherapies [6,7]. As such, building up 3D cellular organizations has become a key target in providing biologically relevant *in vitro* systems for cancer research [8]. In this context, many research efforts have been dedicated to the engineering of spheroid models using various strategies such as hanging drop method [9,10], rotating walls culture [11], hydrogels [12-17], or non-adherent substrates including bovine serum albumin (BSA) [18,19], polyHEMA [20,21], agarose [5,22,23] or Pluronic F [24,25].

Beside the difficulties for obtaining stable spheroids with controlled homogeneity, two key challenges can be emphasized for the development of *in vitro* cancer models. The first one is the ability to tune the number of cells, and the size of the spheroids so as to reproduce metastasis evolution during different stages of ovarian cancer, as well as different types of cancer. A second important challenge is to recapitulate the multidimensional and constantly evolving tumor microenvironment, which also includes the structural matrix, shear stress and soluble – circulating – factors. To address all of these, we first used micro-structured supports – called patches – with repellant coating for the fabrication of size-controlled spheroids with high efficiency (Figure 1A,B). Here, we show that by playing with the dimensions of the patch, we can vary the number of cells in and the size of the spheroids, and reproducibly obtain stable OC spheroids that can easily be handled for further use. Next, spheroids were placed on a second type of patch coated with a thin membrane of type I collagen mimicking the extracellular matrix (ECM-patch) to observe substrate dependent spheroid disruption (Figure 1C,D). This approach should be spheroid-type independent and applicable to assays under different experimental conditions. Of note, the self-standing area of the ECM membrane device has a low stiffness (< 20 kPa) and a large permeability, which play important roles and is unique compared to both conventional culture substrates and plastic porous membranes. As an illustration, this device effectively allows to monitor the invasive behavior of SKOV-3 human OC cells providing a unique insight into cells transitioning within the epithelial to mesenchymal transition spectrum.

**Figure 1.**
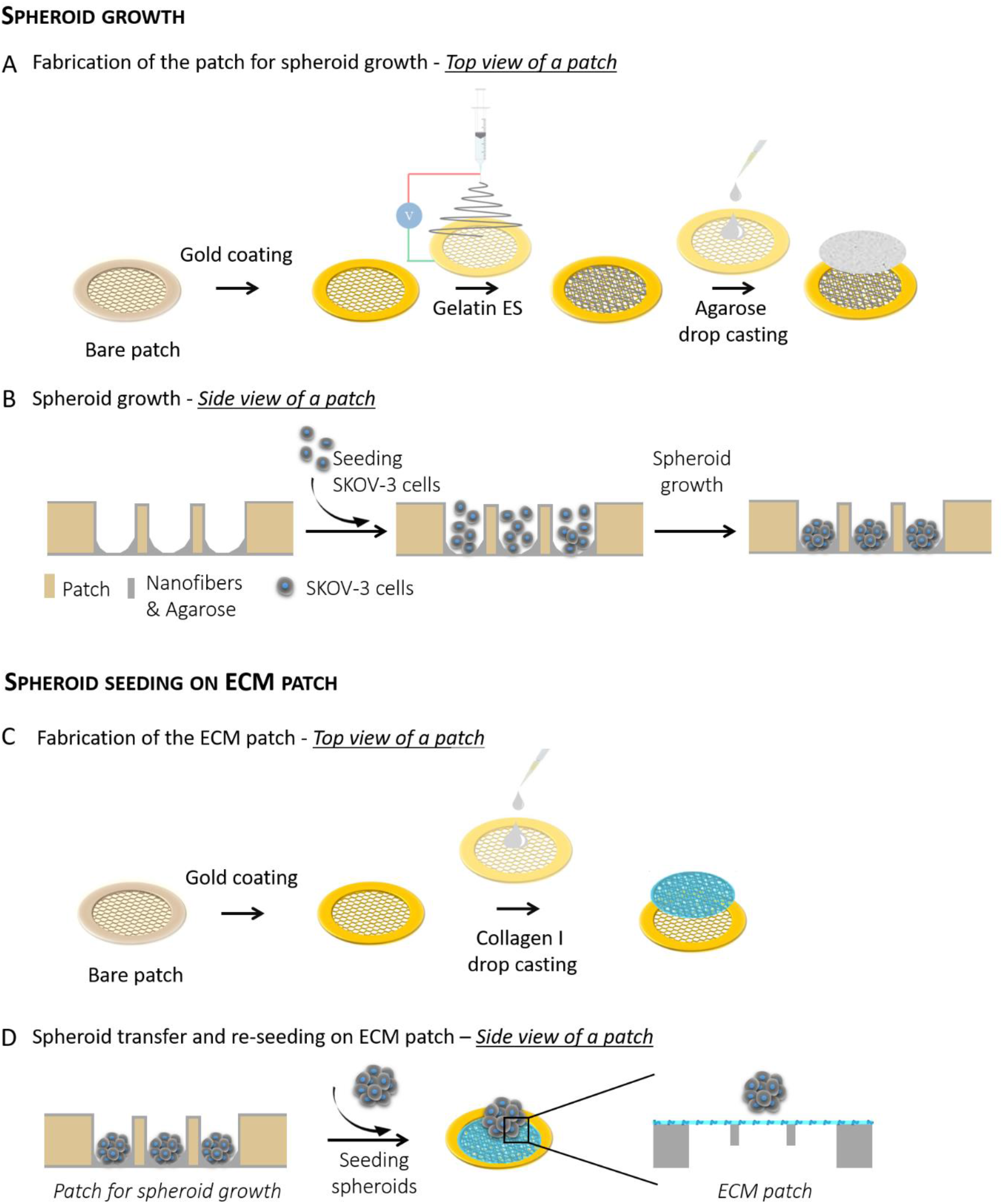
Schematic diagram of micro device fabrication for tumor cell growth and processing. (A) Fabrication of the gelatin/agarose patch for spheroid growth. (B) Tumor spheroid formation in honeycomb compartment with an agarose membrane. (C) Fabrication of the ECM patch and (D) spheroid plating on ECM patch.

## 2. Materials and methods

### 2.1 Fabrication of the patches

The patches were fabricated by photolithography and soft-lithography. SU-8 molds were fabricated by photolithography. For the 50 μm-high patch, SU-8 3050 was spin coated on a silicon wafer at 3 000 rpm for 30 s. For the 200 μm-high patch, SU-8 3050 was spin coated on a silicon wafer at 2 000 rpm for 30 s twice, followed each time by a soft bake. Then, a honeycomb frame of 200 or 400 μm in width was directly patterned by UV exposure at 250 mJ/cm². This SU-8 mold was then exposed in trimethylchlorosilane (TMCS, Sigma, France) vapor for 10 min for surface treatment. Afterward, a mixture of PDMS (GE RTV 615) pre-polymer and its cross-linker at a weight ratio of 10:1 was casted on the SU-8 mold. After curing at 75°C for 4 hours, the PDMS layer was peeled off. Then, this replicated PDMS structure was placed on a glass plate and a solution of a photo-crosslinking polymer (Ormostamp®) was injected in the free space of the PDMS-glass assembly, followed by UV exposure at 1500 mJ/cm², and removal of the PDMS mold. The patches were then coated with gold by sputter deposition for electrospinning (for the fabrication of the spheroid-patches, Figure 1A,B) or for improving hydrophilicity and drop casting of type I collagen (for the fabrication of the ECM patch, Figure 1C,D).

### 2.2 Electrospinning of gelatin

Electrospinning is the extrusion of a soluble biopolymer assisted with electrical field. It is interesting for cell biologists because it provides a net-like structure made of cross-linked nanofibers mimicking the ECM [26]. The gelatin solution was prepared by dissolving 100 mg of gelatin in a mixture of 420 μL acetic acid, 280 μL ethyl acetate and 200 μL DI water. Citric acid (10 mg) was then added as thermal cross-linking agent before electrospinning. The mixture was stirred for 4 hours at room temperature. Gelatin was electrospun onto a gold-coated patch fixed on a tin foil (7 cm in diameter) at a voltage of 11 kV for 7 minutes at a flow rate of 0.2 mL/h, with a controlled humidity of 35% at room temperature (20∼25°C). The distance between the metal needle and the patch was 9-12 cm. After electrospinning, the patch was detached from the tin foil and transferred into an oven working at 140°C for 4 hours for high temperature cross-linking of gelatin.

### 2.3 Agarose drop casting

To promote the formation of spheroids, the gelatin nanofiber layer was coated with a solution of agarose (Fisher Scientific, France). 20 μL of 0.2 w/v% solution in DI water was poured on the patch and let dry in air at room temperature. Note that the gelatin fiber layer was necessary to maintain the agarose film on the porous patch, while preserving the porosity of the support.

### 2.4 ECM patch

Type I collagen was extracted and purified from rat tail tendons as previously described by substituting 500 mM acetic acid with 3 mM hydrochloric acid [27,28]. Collagen purity was assessed by electrophoresis and its concentration estimated by hydroxyproline titration [29]. Type I collagen (0.5 mg/mL in PBS, pH=9.0) was drop cast by pouring 20 μL of the solution on a patch with dimensions: 50 μm thickness and 400 μm width of honeycombs and let dry in air at room temperature.

### 2.5 Second harmonic generation / 2-photon excited fluorescence

We used a custom-built laser-scanning multiphoton microscope and recorded Second Harmonic Generation (SHG) images as previously described [30]. Excitation was provided by a femtosecond titanium–sapphire laser (Mai-Tai, Spectra-Physics) tuned to 860 nm, scanned in the XY directions using galvanometric mirrors and focused using a 25× objective lens (XLPLN25XWMP2, Olympus), with a resolution of 0.35 μm (lateral) × 1.2 μm (axial) and a Z-step of 0.5 μm for the acquisition of Z-stack images. We used circular polarization in order to image all structures independently of their orientation in the image plane, using 100 kHz acquisition rate and 420×420 nm^2^ pixel size.

### 2.6 SEM imaging

Samples were coated with gold for 60 s with a sputtering current of 50 mA before imaging. Samples were fixed on conductive-tapes for imaging with a TM3030 Tabletop Microscope (Hitachi High-Technologies Corporation, Japan) equipped with TM3030 software and working at an acceleration voltage of 15 kV.

### 2.7 Cell culture

SKOV-3 cells (ATCC1, HTB77™), human ovarian adenocarcinoma cell line, were purchased from ATCC (American Type Culture Collection, Manassas, VA). SKOV-3 cells were cultured in RPMI-1640 glutaMAX containing 0.07% sodium bicarbonate supplemented with 10% fetal calf serum and 100 U/mL penicillin streptomycin (all reagents were purchased from Thermo Fisher Scientific). Cells were cultured in T25 cell culture flasks in a humidified air atmosphere with 5% CO_2_ at 37 °C.

### 2.8 Invasion assay

ECM patches were immersed in 70% ethanol for 5 min for sterilization, and washed 3 times for 5 min in sterile PBS. SKOV-3 spheroids, 3 days old, were recovered and seeded on ECM-patch. Patches were then transferred to a controlled atmosphere (37 °C, 5% CO_2_) before immunostaining of the cells.

### 2.9 Immunofluorescence staining

Cells were fixed in 4% paraformaldehyde (PFA) in PBS for 10 minutes, rinsed three times with PBS. The cells were permeabilized for 4 minutes with 0.1% Triton x100 in PBS at 4 °C, washed again and saturated with PBS containing 0.5% BSA for 30 min. For spheroid staining, cell junctions were stained overnight at 4 °C with a primary E-cadherin (610181, BD Biosciences) staining and secondary IgG, IgM (H+L) Goat anti-Mouse, Alexa Fluor™ 488, Invitrogen™ for 1 hour at room temperature. Cells were washed 3 times with PBS for 5 min each time to remove unbound antibody. For additional staining, we used EPCAM CD326 monoclonal antibody conjugated with Alexa Fluor® 488 (eBioscience™). Tumor cells were incubated overnight at 4 °C at a 1/20 dilution and washed 3 times with PBS for 5 min each time to remove unbound antibody.

For the invasion assay, cells were incubated overnight at 4 °C with recombinant anti-vimentin antibody conjugated with Alexa Fluor® 568 (ab202504, Abcam®) at a 1/1000 dilution. After washing the actin cytoskeleton was stained with Alexa Fluor® 488 Phalloidin in PBS (containing 1% DMSO from the original stock solution, Abcam®) for 40 min at room temperature in a dark chamber. Cell nuclei were then stained with DAPI (4,6-diamidino-2-phenylindole dihydrochloride, Molecular Probe®) for 15 min.

Immunofluorescent labelling was observed with a confocal microscope (LSM710, Zeiss) equipped with 405, 488, and 543 nm lasers and with LSM ZEN 2009 software and using 1 μm z-stack intervals and sequential scanning. Images were processed with ImageJ.

## 3. Results and discussion

### 3.1 Patch device and coating

Figure 1A-B describe our strategy for spheroid growth and control of the number of cells and spheroid size. We use microfabricated *patches* that are circular culture supports, 1.3 cm in diameter, with a honeycomb microporous pattern. These patches were fabricated by photolithography and soft-lithography for obtaining a PDMS replica. Then the patch can be obtained by injecting a photo-crosslinking polymer to the replica on a glass, and UV-assisted curing. The porosity of the microframe presents many advantages notably for limiting the contribution of the mechanical properties of a solid substrate, while preserving the biological permeability for exchanges of circulating soluble factors. These supports have been shown to allow the maintenance and differentiation of human induced pluripotent stem cells (hiPSCs) over days [31-37]. Patches were coated with gold prior to electrospinning of gelatin (Figure 1A). On top of the electrospun gelatin layer, agarose is drop cast ensuring a non-adherent coating. After this step, human ovarian carcinoma – SKOV-3 – cells were seeded and allowed to form spheroids within the honeycomb chambers (Figure 1B).

One important challenge in the field is to control the size and homogeneity of the spheroids. To this aim, we have changed the dimensions of the honeycomb microframe: the width (W) of the honeycomb (200 and 400 μm) and the height (H) of the walls (50 and 200 μm) (Figure 2A). Two patches were fabricated: W200H50 and W400H200 (Figure S1).

**Figure 2.**
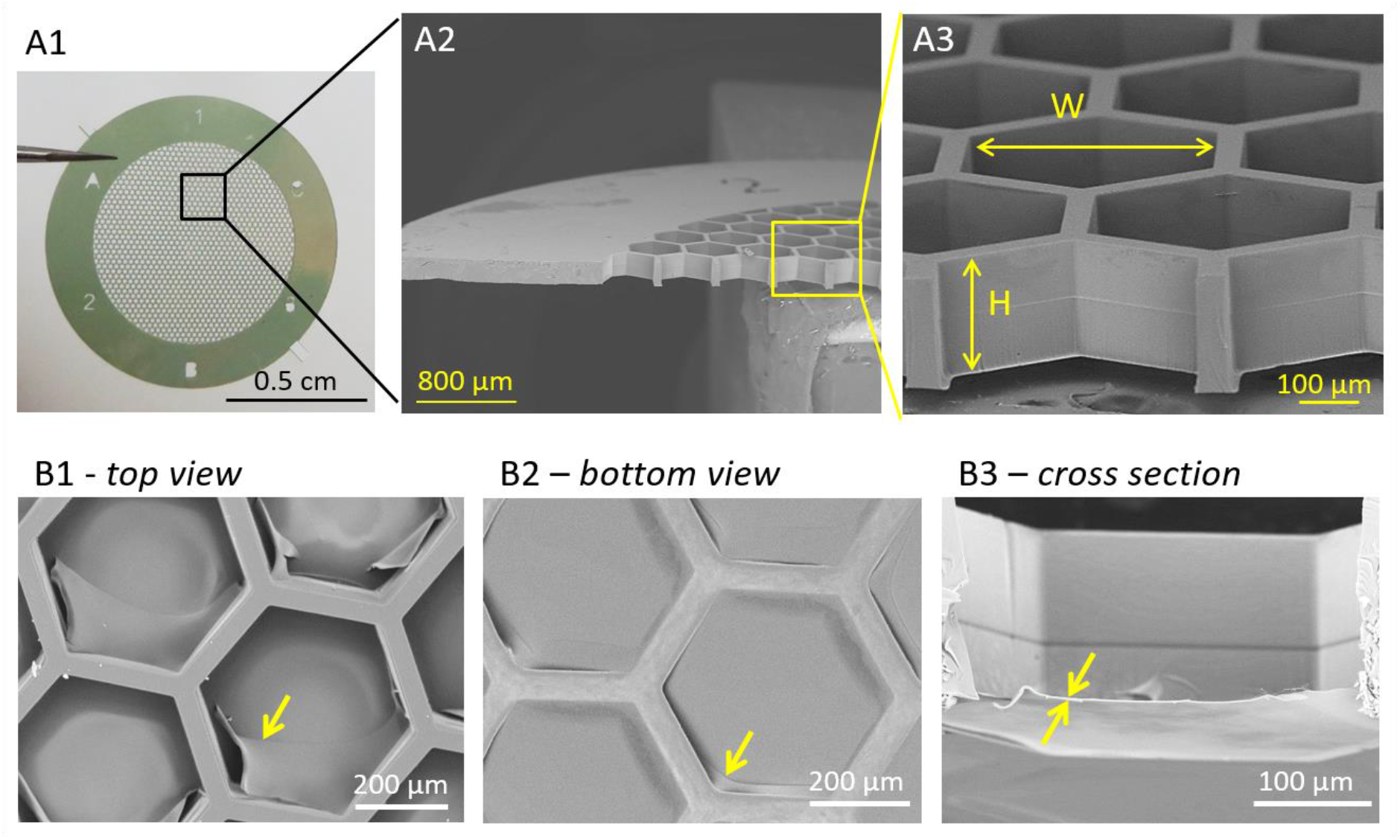
(A) Bare patch: (A1) Photograph and (A2,3) SEM images of the fabricated W400H200 patch. (B) SEM images of the coating of patches with gelatin nanofibers topped with agarose film (see yellow arrows) for spheroid growth: (B1) top view, (B2) bottom view and (B3) cross-section.

Gelatin was successfully electrospun and crosslinked giving rise to a homogeneous layer of nanofibers whatever the dimensions of the microframe (Figure S2). A solution of agarose was subsequently drop cast, giving rise to a homogeneous non-adherent film that was successfully maintained in 200 μm high honeycombs (Figure 2B).

### 3.2 Growth of human ovarian spheroids

After coating the patch, SKOV-3 cells have been seeded on the W200H50 and W400H200 patches. Different cell seeding densities have been tested. We have selected 100 000 cells per patch and 400 000 cells per patch for W200H50 and W400H200 respectively that were found to be the maximal cell density allowing to grow a single spheroid per honeycomb compartment and preventing overflow of cells into layers (Figure S3). Figure 3A shows the formation of the spheroids during the first three days after seeding SKOV-3 on W200H50 patches. Starting from isolated cells, at day 0, we can clearly see the aggregation of cells into circular structures that occurs after one day and slowly evolves going from 91 μm to 109 μm in diameter after three days (Figure 3B and Table S1). The spheroids obtained after three days exhibit an aspect ratio of 1 (Figure S4) with a Gaussian distribution of their size (Figure 3C). The same trend is observed after seeding cells on W400H200 patches, going from isolated to rapidly aggregating assemblies obtained after one day (Figure 3D). These assemblies grow slowly, going from 214 to 235 μm after three days (Figure 3E and Table S2). After three days, homogeneous spheroids with an aspect ratio of 1 were obtained (Figure 3F and Figure S5). Interestingly, the spheroids are more than twice as large in diameter as those obtained with the W200H50 patches. Importantly, in both systems, nearly all honeycombs contained a spheroid. Given that W200H50 and W400H200 are made up of *ca.* 1350 and 400 honeycombs respectively, we expect a minimum of *ca.* 70 and 1000 cells per spheroid with respect to the initial cell density.

**Figure 3.**
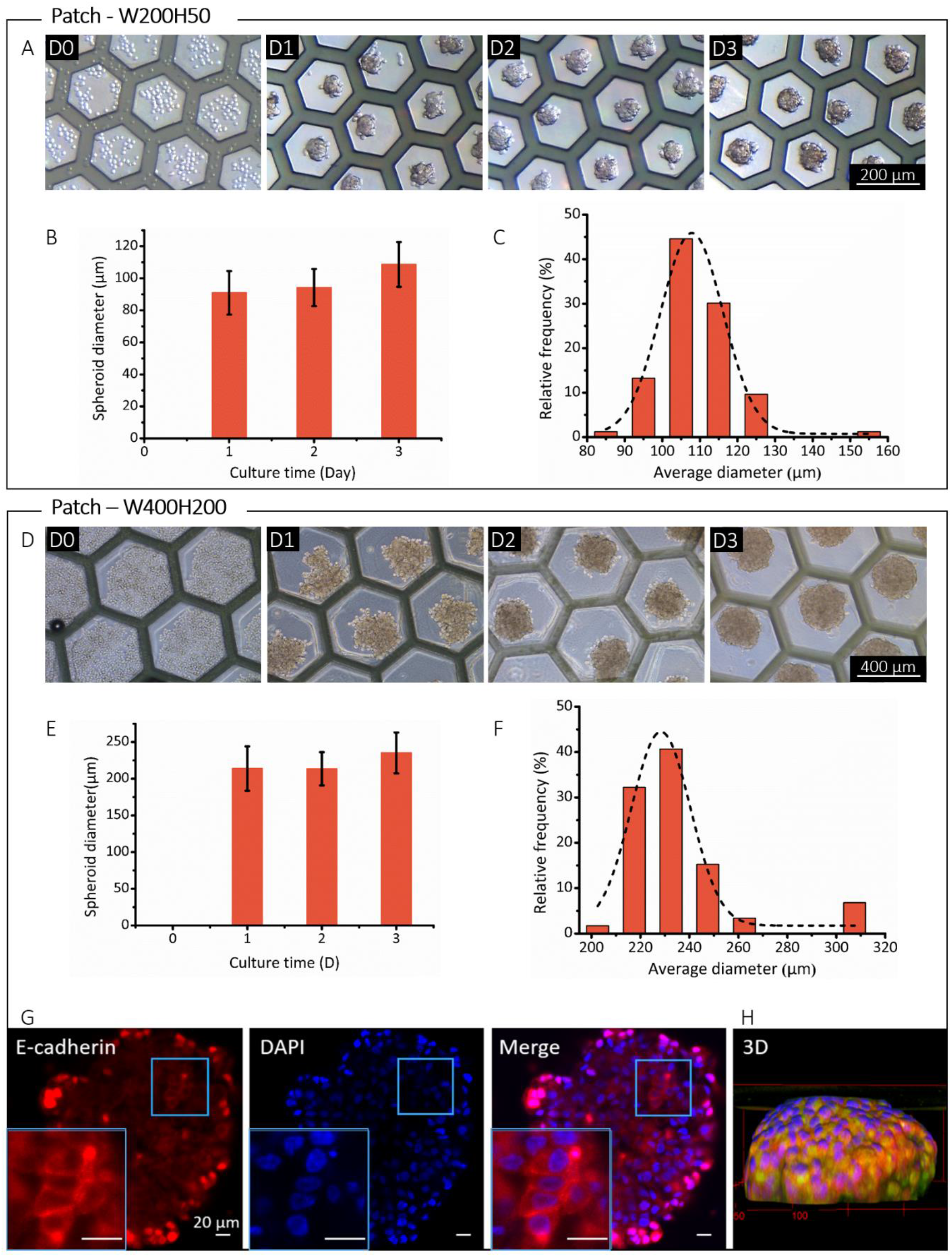
Spheroid growth on (A-C) W200H50 and (D-G) W400H200 patches. (A,D) Optical microscopy images of spheroid growth, 1, 2, and 3 days after SKOV-3 seeding. (B,E) Histograms of the spheroid diameters as a function of culture time, and (C,F) distribution of spheroid size 3 days after SKOV-3 seeding. (G) Immunofluorescence image of a SKOV-3 spheroid (Z-slice) and (H) 3D reconstruction of a second spheroid (epithelial markers EpCAM (in green) and E-cadherin (in red) and nuclei (DAPI in blue).

The 3D spherical aspect of the spheroids was further checked using immunofluorescence microscopy (Figure 3G). Staining was directed towards E-Cadherin cell junctions, which is an epithelial marker, together with the staining of the nuclei (DAPI staining). These observations show that the fluorescence intensity of both markers is higher at the edge of the spheroid compared to the core due to the limited diffusion of the fluorescent probe within the dense cellular aggregate. However, E-cadherin cell junctions are clearly observed (Figure 3G insert). In parallel, spheroids were stained for vimentin, a cytoskeletal protein upregulated during cell transitioning events and whose expression is associated with mesenchymal phenotypes, showing a hardly detectable signal (Figure S6). These observations highlight the specificity of the epithelial marker staining and indicate the epithelial phenotype of the SKOV-3 cells within the spheroids. Very importantly, the 3D reconstruction confirmed the spherical, slightly flattened, aspect of the spheroid.

### 3.3 Invasion assay: inducing epithelial-to-mesenchymal transition

Tumor spheroids obtained were stable and could be handled and removed easily to be re-seeded on a second type of patches mimicking the ECM. These biologically relevant ECM patches are obtained by drop casting of type I collagen as model of connective tissues (Figure 1C,D). Figure 4A shows the characterization of the ECM patch using multiphoton microscopy based on second harmonic generation (SHG) [30]. This imaging modality allows to monitor unlabeled collagen I with high specificity towards its fibrillary hierarchical organization. These images show the fibrillary structure of the network, which is few micron in thickness. After seeding of the spheroids, migration could be observed with highly different phenotypes from the core to the front of the spheroidal aggregate (Figure 4B). Staining of the nucleus (DAPI staining) and actin (phalloidin staining) were combined with the immunostaining of vimentin, which is commonly used as a marker of epithelial-to-mesenchymal plasticity. Fluorescence microscopy images revealed the cellular heterogeneity associated to the invasive migration of metastatic cell aggregates. In the core, epithelial phenotypes could be observed, as previously identified within the spheroids (Figure 4C1). At the front of the aggregate, cells with different phenotypes could be observed, characterized by elongated nucleus and a high cortical expression of vimentin, which co-localized with the actin cytoskeleton (Figure 4C2). This is characteristic of mesenchymal phenotypes and shows that cells transitioning within the epithelial-to-mesenchymal spectrum in the course of the invasion process is reproduced on the patch.

**Figure 4.**
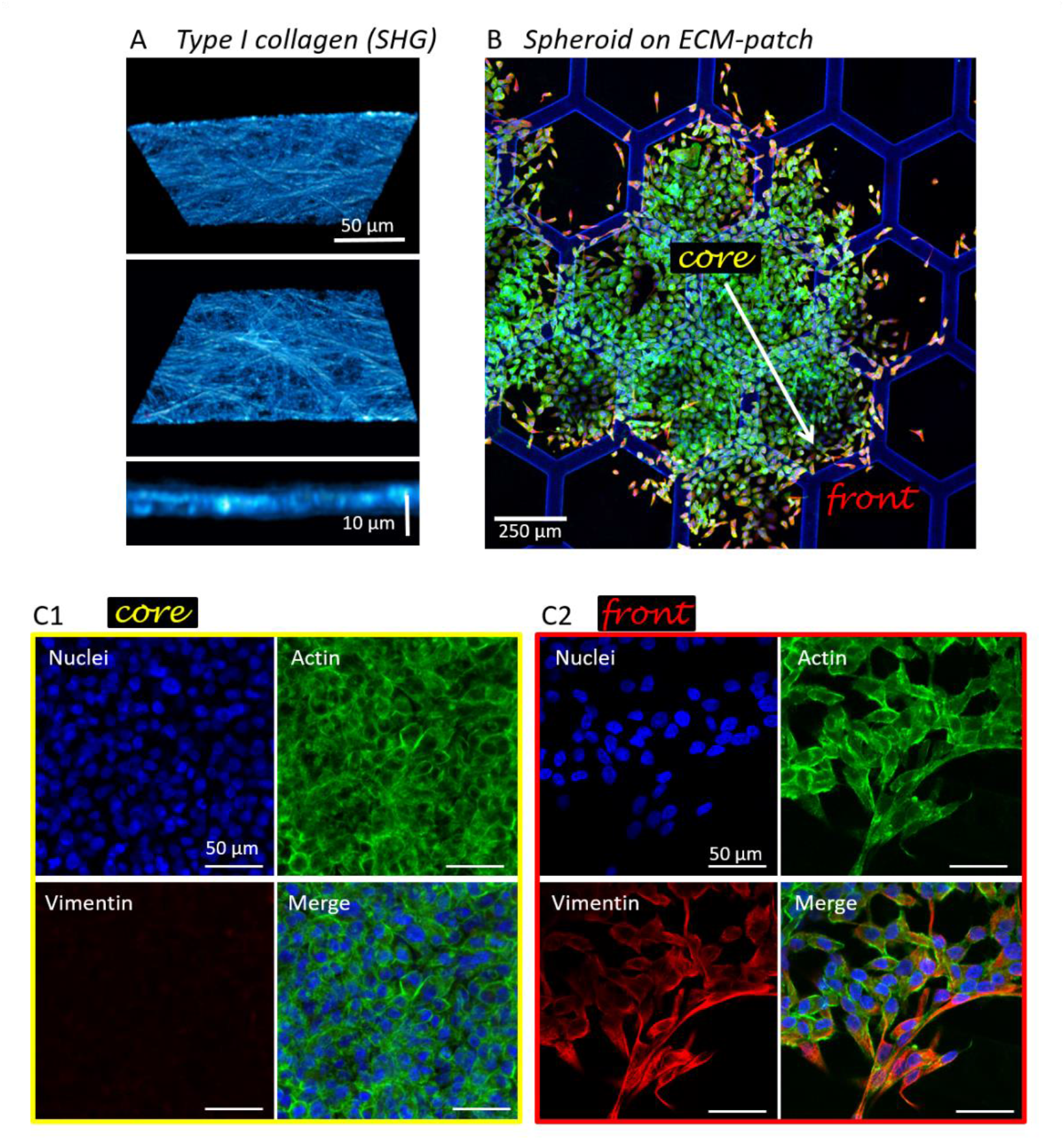
(A) Top, bottom and side view of the ECM patch using SHG. (B-C) Immunofluorescence image of SKOV-3 spheroids from W200H50 patches two days after seeding on ECM-coated patch. Cells were stained for vimentin (red), actin (green), and nuclei (blue).

Finally, one important advantage of using this strategy is the possibility to integrate the patch into a diversity of microfluidic chips for the development of various bioassays [24,34-37]. This will be particularly relevant for modeling ovarian cancers that are very sensitive to their surrounding microenvironment [38], which includes the soluble environment, the ascites that is an excess fluid resulting from lymphatic obstruction found in physio-pathological conditions [39] and that has been shown to play a role in EMT [40].

## Conclusions

We have developed a robust strategy to control the size and the number of cells in spheroids obtained from human OC cells. Playing with the dimensions of porous microfabricated supports allowed to grow spheroids from SKOV-3 cells with diameter in the 90 to 235 μm range. We have shown that spheroid growth can be achieved over a short period of time (1 to 3 days) and with high-throughput (from 400 to more than 1300 spheroids depending on the patch). These stable spheroids can be handled and transferred on different kinds of supports. This provides a new *in vitro* model, which recapitulates the tumor environment with varying metastatic assemblies and adjustable microenvironment structure. Because these patches can be easily incorporated within microfluidic chips, it provides a mean to control and explore independently the impact of the shear stress and soluble factors (flow and circulating environment) on metastatic behaviors of OC cells. Hence, this strategy is an asset to adapt genetic tools to tune collective and single cell behavior as a function of the microenvironment, in a flexible and easy way. The invasion assay will provide the biological and biomedical community with unique tools for investigating collective migratory behaviors and epithelial-to-mesenchymal transition at play in physiological and pathological context typically embryogenesis and cancer. The possibility to independently control the metastasis size, microenvironment structure and screen anti-cancer drugs represents a new perspective for cancer-on-a-chip research.

## Acknowledgements

This project has received financial support from the CNRS through the. Changchong Chen thanks the China Scholarship Council for his PhD grant.

