## Supplementary materials for "High-throughput tuning of ovarian cancer spheroids for on-chip invasion assays"

### Supporting Information

**1. Microfabrication and coating of the honeycomb patches.**

**2. Varying cell seeding density on W200H50 patches.**

**3. Spheroids obtained from W200H50 patches: size and aspect ratios.**

**4. Spheroids obtained from W400H200 patches: size and aspect ratios.**

**5. Vimentin staining of the spheroids.**

#### 1. Microfabrication and coating of the honeycomb patches.

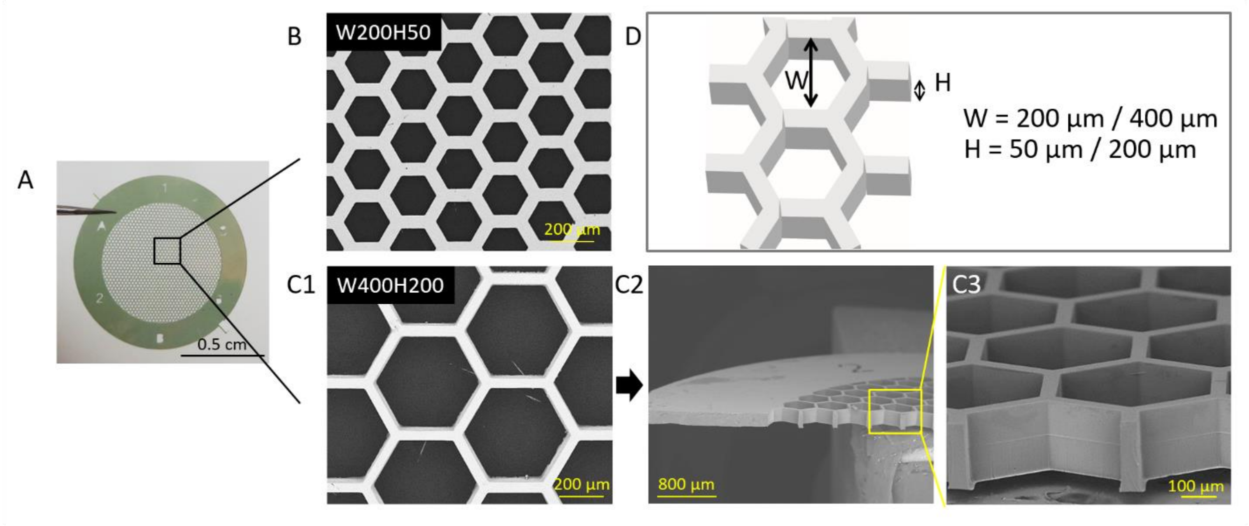


**Figure S1.** (A) Photograph and (B-C) SEM images of the microfabricated culture support - called *patch* - with varying width and thickness of the honeycomb walls: (B) W200H50 patch, and (C1-C3) W400H200 patch. (D) Scheme of the honeycomb and varied dimensions.

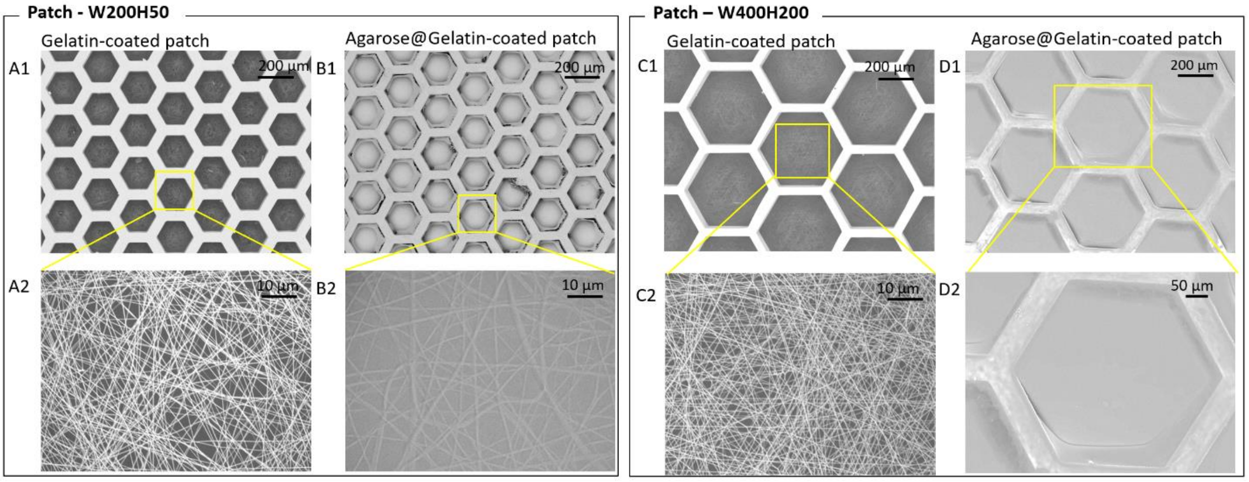


**Figure S2.** SEM images of the coating of patches for spheroid growth: (A-B) W200H50 and (C-D) W400H200 patches coated with (A-C) gelatin nanofibers and (B-D) agarose.

#### 2. Varying cell seeding density on W200H50 patches.

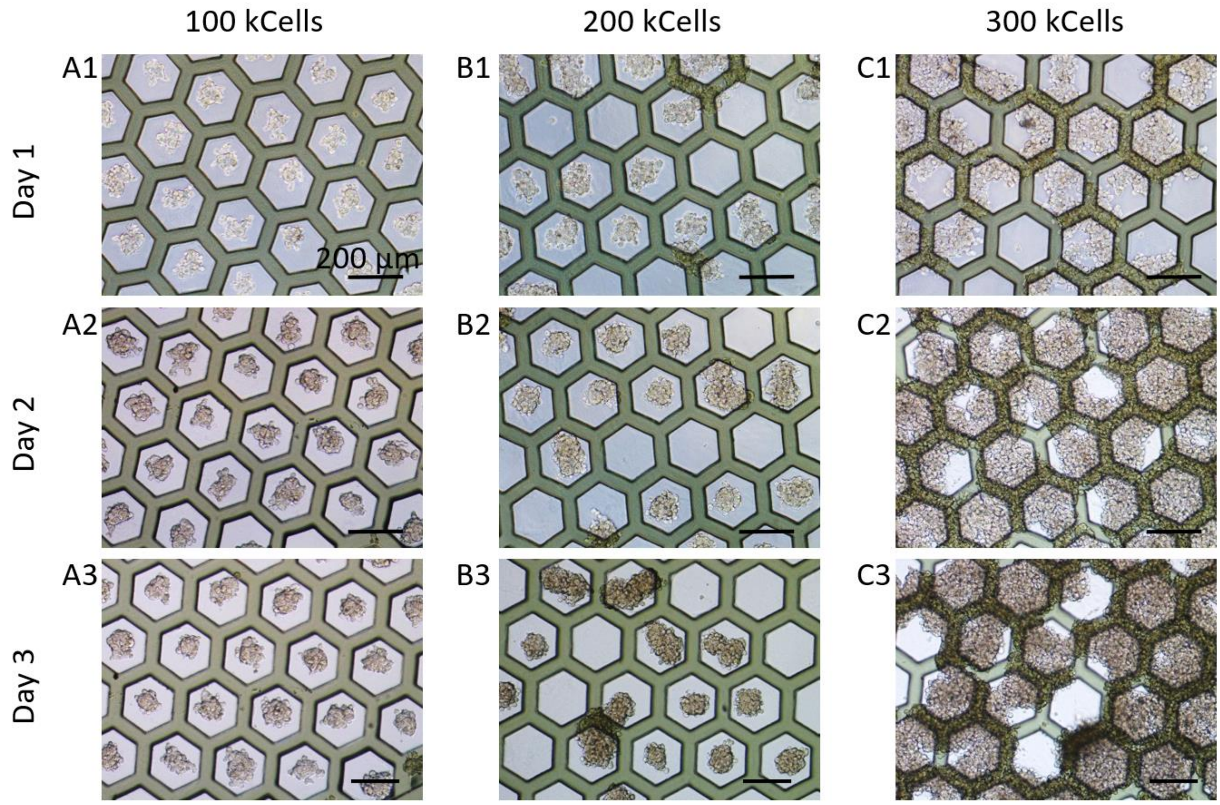


**Figure S3.** Optical microscopy images of cellular aggregates obtained after seeding SKOV-3 cells onto W200H50 patches after 1, 2 and 3 days of culture and at different cell seeding densities: (A) 100 000, (B) 200 000, (C) 300 000 cells.

#### 3. Spheroids obtained from W200H50 patches: size and aspect ratios.

| Culture time (day) | Size (µm) | Standard deviation (µm) |
|---|---|---|
| 1 | 91 | 14 |
| 2 | 94 | 12 |
| 3 | 109 | 14 |

**Table S1.** Size of the spheroids obtained 1, 2, or 3 days after seeding SKOV-3 cells on W200H50 patches.

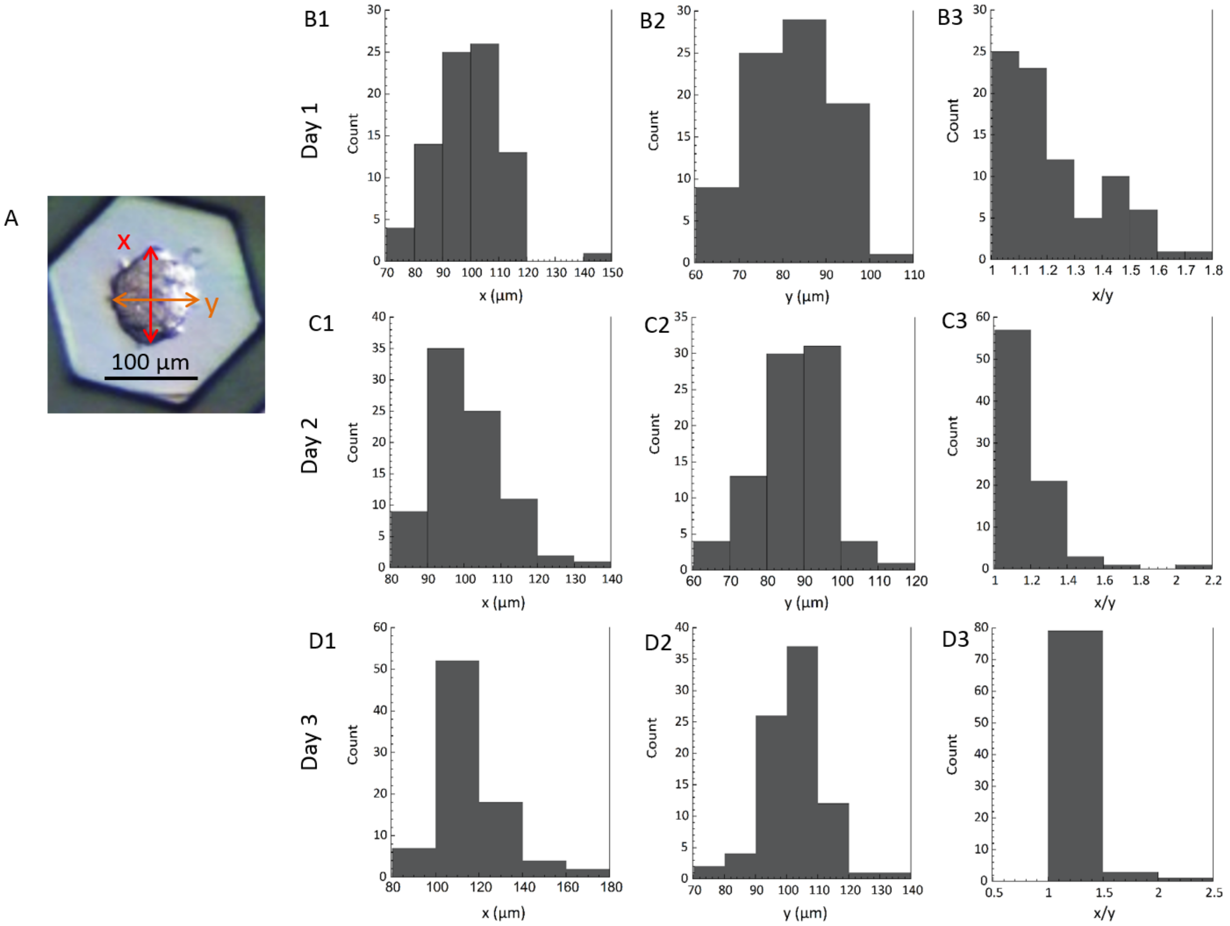


**Figure S4.** Distribution of the measurements of the two dimensions – (1) x and (2) y - and (3) resulting x/y aspect ratio of the spheroids obtained after seeding SKOV-3 cells on W200H50 patches. (A) Typical optical microscopy image; and measurements of the spheroids obtained after (B) one, (C) two, and (D) three days of culture.

#### 4. Spheroids obtained from W400H200 patches: size and aspect ratios.

| Culture time (day) | Size (µm) | Standard deviation (µm) |
|---|---|---|
| 1 | 214 | 30 |
| 2 | 213 | 23 |
| 3 | 235 | 28 |

**Table S2.** Size of the spheroids obtained 1, 2, or 3 days after seeding SKOV-3 cells on W400H200 patches.

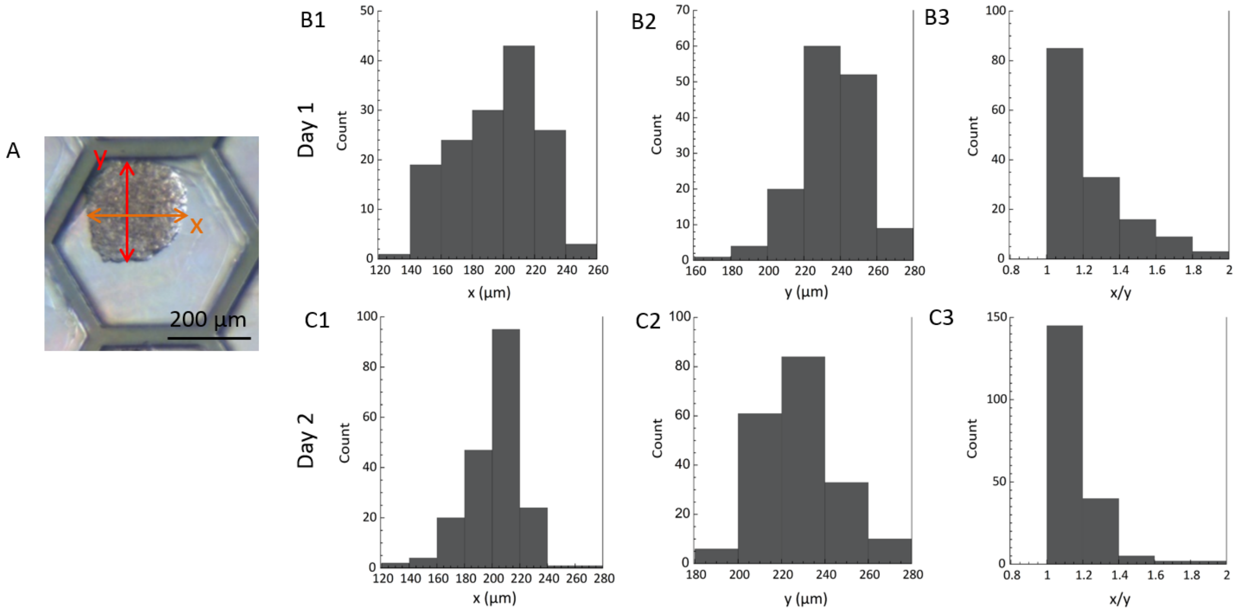


**Figure S5.** Distribution of the measurements of the two dimensions – (1) x and (2) y - and (3) resulting x/y aspect ratio of the spheroids obtained after seeding SKOV-3 cells on W400H200 patches. (A) Typical optical microscopy image; and measurements of the spheroids obtained after (B) one and (C) two days of culture.

#### 5. Vimentin staining of the spheroids.

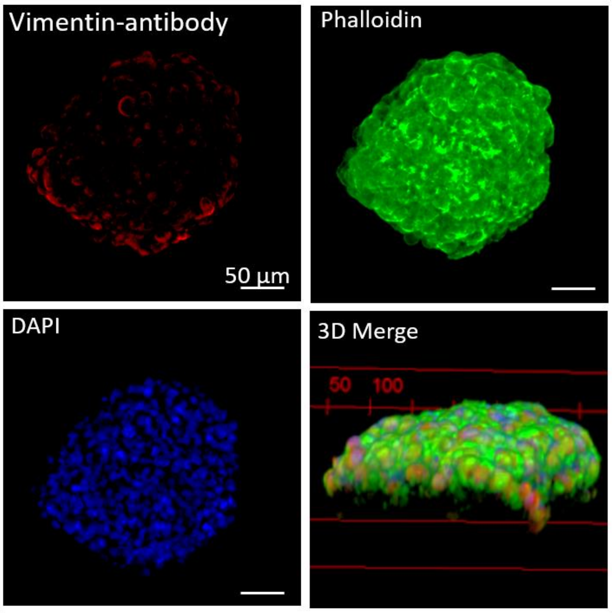


**Figure S6.** Immunofluorescence image of SKOV-3 spheroids (Z-slice) and 3D reconstruction.
